# Love-thy-neighbor: Neural networks for tracking and lineage tracing in budding yeast

**DOI:** 10.64898/2026.01.09.698579

**Authors:** Marija Zelic, Vojislav Gligorovski, Farzaneh Labbaf, Marco Labagnara, Roxane Oesterle, Greta Brenna, Fanny Massard, Samir G. Chethan, Wanlan Li, Sophie G. Martin, Silke Hauf, Sahand Jamal Rahi

## Abstract

Tracking and lineage tracing are widely needed tasks in biological image analysis. For cells that grow and divide, tracking is challenging because cells change in number, shape, and size throughout a recording. As the time interval between images increases, it becomes more difficult to establish correspondences between cells across timepoints. Consequently, tracking has to be performed between consecutive or temporally close images, which leads to exponentially decreasing tracking accuracy and thus high sensitivity to error rates. For budding yeast, this challenge is further heightened by the similarity of cells in colonies, their dense packing, the asymmetric nature of cell divisions, and movement due to growth of the colony. A related task, lineage tracing, is similarly challenging without fluorescent markers due to multiple potential mother cells surrounding a new daughter cell. Here, we present neural networks for budding yeast tracking and lineage tracing, named LYN-track and LYN-trace, respectively. These methods leverage fine geometric features of cells and their neighborhoods. To train and test the algorithms, we recorded and annotated new budding and fission yeast microscopy movies (78,852 frame-to-frame tracklets, 2,512 images), which we make freely available. On these and existing datasets, our neural network-based methods demonstrate robust, above state-of-the-art performance. Both tools have been integrated into graphical user interfaces (GUIs), available on Github, and can be straightforwardly retrained with custom data if desired.

## Introduction

Labeling cells consistently across time (tracking) and identifying the mother cell to a new daughter cell (lineage tracing) are two widely shared needs in processing microscopy images. Tracking and lineage tracing are particulary difficult when cells are densely packed, move, change in number and shape, and are poor in distinguishing features, which is the case for typical budding yeast timelapse recordings^1–7^. Budding yeast mothers give rise to a smaller daughter once per cell cycle, while fission yeast divides symmetrically into two daughter cells. These features require tracking to be performed between images that are close in time, e.g., from one image to the next, which renders the result highly sensitive to the error rate as incorrect assignments propagate exponentially. (The fraction of correctly tracked cells decreases as ∼ *c^n^* after *n* timepoints if 1 − *c* is the frame-to-frame error rate, which depends on the frame rate.) Thus, lowering the error rate substantially reduces the amount of manual correction needed.

Tracking generally involves three steps post image segmentation^8–17^: First, quantitative characteristics are computed for each cell in each frame, e.g., size and position. Then, pairs of cells from consecutive timepoints are compared to quantify the possibility of being the same cell, e.g., by similarity of size and position. Finally, the pair-wise assignment scores are globally optimized. Existing tracking methods tend to differ more in the first two steps, the quantitative characterization of cells and of their assignments, while the last step is usually performed by solving the linear assignment problem. For budding yeast, a number of tracking algorithms have been devised that do not rely on neural networks.^18–24^ For example, the tracking algorithm in YeaZ^18^ describes each cell by its center-of-mass coordinates and pixel area, inspired by single-particle tracking^25^, which similarly uses position and pixel intensity. The potential correspondence between cells at two consecutive timepoints is quantified by the Euclidean distance between vectors in this feature space. TracX^20^, available as Matlab code, uses a compressed, short-wavelength Fourier representation of each cell image and computes the similarity between cells by the Euclidean distance of the transformed images. FIEST^24^ takes the pixel region of each cell and optimizes for maximal overlap between timepoints.

Existing methods for budding yeast tracking leave a substantial amount of information about cells and their neighborhoods unused. Considering the exponential nature of the decrease in the number of correctly tracked cells in a recording, it is highly desirable to lower frame-to-frame tracking errors. Here, we leverage the hitherto unused information about cell geometries and neighborhoods as well as deep learning for budding yeast tracking. We developed a specially designed graph neural network (GNN)^26^, which uses the geometric features of each cell and its neighbors. In contrast to most previous GNNs for tracking objects such as humans, cars, or mammalian blast cells that take advantage of clearly distinguishing features^27–29^, our novel architecture employs message passing within each timeframe in order to accumulate information about cell neighborhoods. The GNN characterizes cells, their neighborhood, and frame-to-frame correspondences in a single unified model. Furthermore, we built the method to only rely on cell masks, not the underlying microscopy images. Considering that highly accurate segmentation methods now exist for budding yeast image analysis^18,19,22–24,30–34^, we can decouple the tracking algorithm from the variability of the underlying images, guaranteeing robustness to changes in the ‘look’ of the images.

Lineage tracing is another important need in timelapse recording analysis. For budding yeast, the most common approach is to assign the nearest cell as the mother to a newly emerged cell^21^. Clearly, for a new cell appearing in the interior of a colony, this method fails frequently. With multiple z-stack images, more information is available for lineage tracing^23^ but it is uncommon to obtain multiple sections at each timepoint, in part due to the proportional increase in the size of the recordings. Here, we leverage the fact that the biological positioning of a new bud is not random and its growth is not isotropic. Therefore, we propose a fully connected deep neural network (FC-NN) for lineage tracing, which is trained to assign the most likely mother to a bud based on cell geometry and growth patterns.

In summary, we present here:

deep learning methods for tracking and lineage tracing in budding yeast, named ‘Love thY Neighbor’ (LYN)-track and -trace,
a novel GNN architecture for tracking,
substantial number of new recordings with ground-truth annotations for developing tracking and lineage tracing methods.

We expect our methods to substantially speed up analyses of yeast microscopy movies.

## Results

### New and existing datasets for training and benchmarking

To create training and test datasets showing sufficient colony growth and number of cell divisions to thoroughly test tracking and lineage tracing methods, we recorded seven budding yeast colonies in medium conducive to rapid growth in microfluidic chips (rows labeled ‘SJR (New)’ in **Table 1**, **Fig. 1 A**, **Methods**). These recordings comprise 1,197 images and capture 1,203 net new cells, mostly from cell divisions as well as a small number of cells flowing in or out from outside the field of view. In these recordings, 52,164 cells had to be tracked from one timepoint to the next, defining single-cell frame-to-frame ‘tracklets’. We further recorded 9 fission yeast colonies comprising 1,315 images, 964 new cells, and 26,688 tracklets (rows labeled ‘SM (New)’ and ‘SH (New)’ in **Table 1**, **Fig. 1 B, C**). We analyzed the recordings with YeaZ, performing image segmentation and initial tracking. Then, we meticulously refined the results manually using the YeaZ GUI. For establishing ground-truth mother-daughter cell lineages in the budding yeast recordings, we utilized a fluorescent bud neck marker. (Such a marker is generally not available in budding yeast recordings, making lineage tracing challenging.) To further test the generalizability of our methods, we draw on external datasets, specifically, TTS-SC7-Sc from ref.^20^ and seven videos from the Yeast Image Toolkit (YIT)^35^ (**Table 1**, **Fig. 1 D**). We re-annotated these recordings also, except we could not establish ground-truth lineages for the YIT datasets since a bud neck marker was not available.

**Figure 1:**
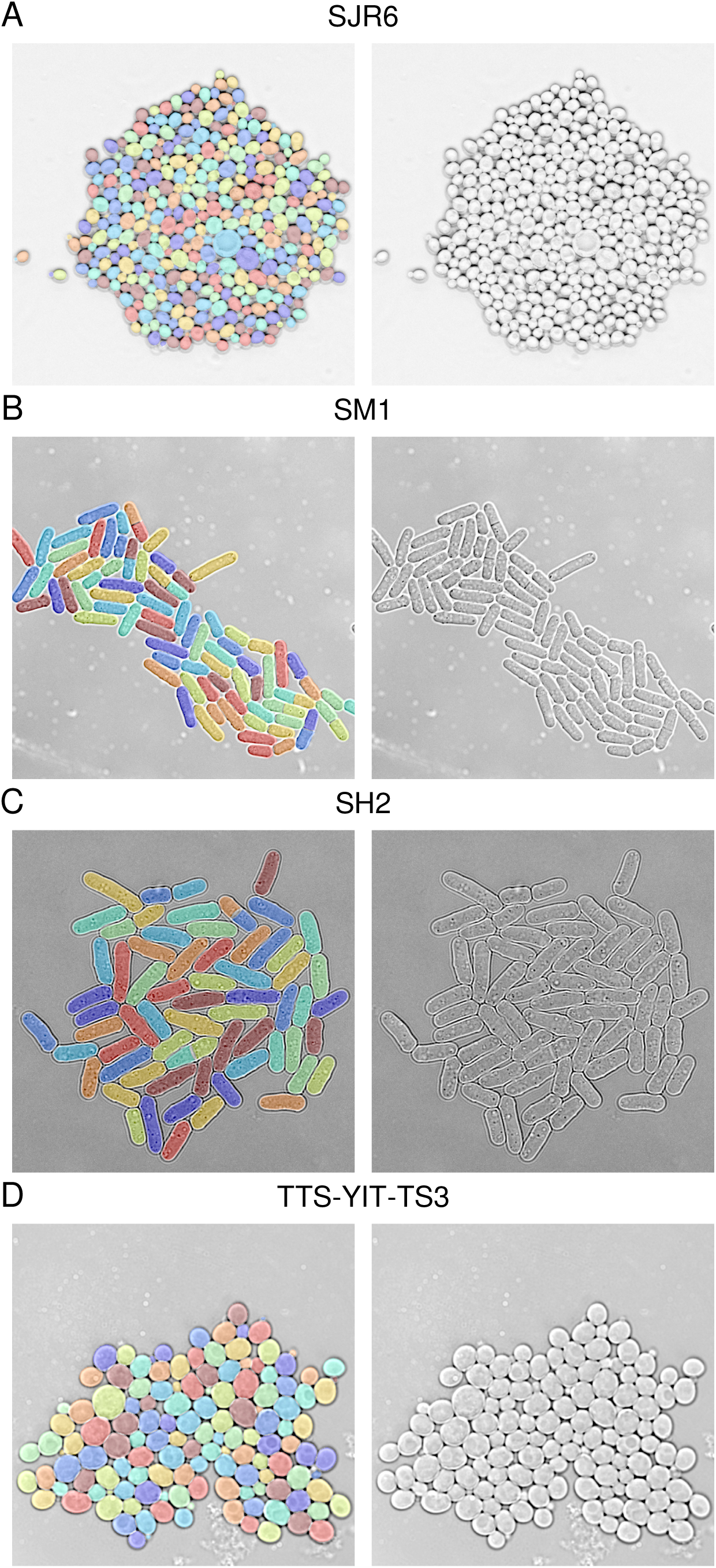
Sample images (right) and segmentation masks (left) from the datasets used in this study. The last image in each timelapse recording is shown.

**Table 1:** Overview of datasets. For budding yeast recordings, ‘# new cells’ describes the net number of cells added to the field of view. In the fission yeast recordings, there was no inflow or outflow of cells, and the number of new cells represents the total number of daughter cells generated after cell division. Tracklets are the number of cells in each frame for which a correspondence needs to be established in the next frame. Sc, *Saccharomyces cerevisiae* (budding yeast). Sp, *Schizosaccharomyces pombe* (fission yeast).

| Source | Name | Species | # frames | # cells at start | # new cells | # tracklets | Frame interval [min] |
| --- | --- | --- | --- | --- | --- | --- | --- |
| SJR (New) | SJR1-Sc | Sc | 181 | 2 | 80 | 5141 | 5 |
|  | SJR2-Sc | Sc | 181 | 7 | 148 | 9805 | 5 |
|  | SJR3-Sc | Sc | 181 | 2 | 94 | 4890 | 5 |
|  | SJR4-Sc | Sc | 181 | 2 | 121 | 5629 | 5 |
|  | SJR5-Sc | Sc | 145 | 3 | 363 | 12042 | 5 |
|  | SJR6-Sc | Sc | 147 | 4 | 361 | 11973 | 5 |
|  | SJR7-Sc | Sc | 181 | 2 | 36 | 2684 | 5 |
| JS<sup>20</sup> | TTS-SC7-Sc | Sc | 120 | 3 | 328 | 7932 | 5 |
| YIT<sup>35</sup> | TTS-YIT-TS1-Sc | Sc | 60 | 14 | 13 | 1123 | 4 |
|  | TTS-YIT-TS2-Sc | Sc | 30 | 4 | 2 | 138 | 3 |
|  | TTS-YIT-TS3-Sc | Sc | 20 | 101 | 33 | 2123 | 3 |
|  | TTS-YIT-TS4-Sc | Sc | 20 | 163 | 62 | 3740 | 3 |
|  | TTS-YIT-TS5-Sc | Sc | 20 | 135 | 20 | 2724 | 3 |
|  | TTS-YIT-TS6-Sc | Sc | 10 | 34 | 15 | 384 | 4 |
|  | TTS-YIT-TS7-Sc | Sc | 10 | 126 | 43 | 1293 | 4 |
| SM (New) | SM1-Sp | Sp | 180 | 2 | 164 | 3743 | 10 |
|  | SM2-Sp | Sp | 180 | 2 | 108 | 2772 | 10 |
|  | SM3-Sp | Sp | 180 | 2 | 152 | 3357 | 10 |
|  | SM4-Sp | Sp | 180 | 1 | 116 | 2398 | 10 |
| SH (New) | SH1-Sp | Sp | 145 | 7 | 118 | 4132 | 5 |
|  | SH2-Sp | Sp | 145 | 6 | 122 | 4439 | 5 |
|  | SH3-Sp | Sp | 75 | 10 | 62 | 1819 | 5 |
|  | SH4-Sp | Sp | 115 | 7 | 92 | 3068 | 5 |
|  | SH5-Sp | Sp | 115 | 2 | 30 | 960 | 5 |

### Overview of GNN-based tracking algorithm LYN-track

Our tracking algorithm begins with segmented images (Steps 1 and 2 in **Fig. 2**) and extracts quantitative features for each cell and for pairs of cells in the same frame (**Table 2**, Step 3 in **Fig. 2**). Subsequently, an input graph is constructed (Step 4): For each pair of cells, one cell from the first frame and the other cell from the second frame, a node is added to the graph, which represents the combined features of both single cells. The nodes are connected by edges (indicated by blue lines in **Fig. 2**), which link four cells: two cells in the same and two cells in the next frame. These edges are associated with the combined features of the two pairs of cells, with each pair being in the same frame. If either pair in the same frame is further separated than a pre-defined distance threshold, the edge is removed (edges that remain indicated by black lines in **Fig. 2**). The GNN is trained to identify the correct nodes in the input graph, i.e., the correct associations of cells between the two frames (Step 5). After training, the GNN scores tracklets by assigning a score to each node. The overall best tracking across two frames is obtained by solving the linear assignment problem, yielding the assignment matrix (Step 6).

**Figure 2:**
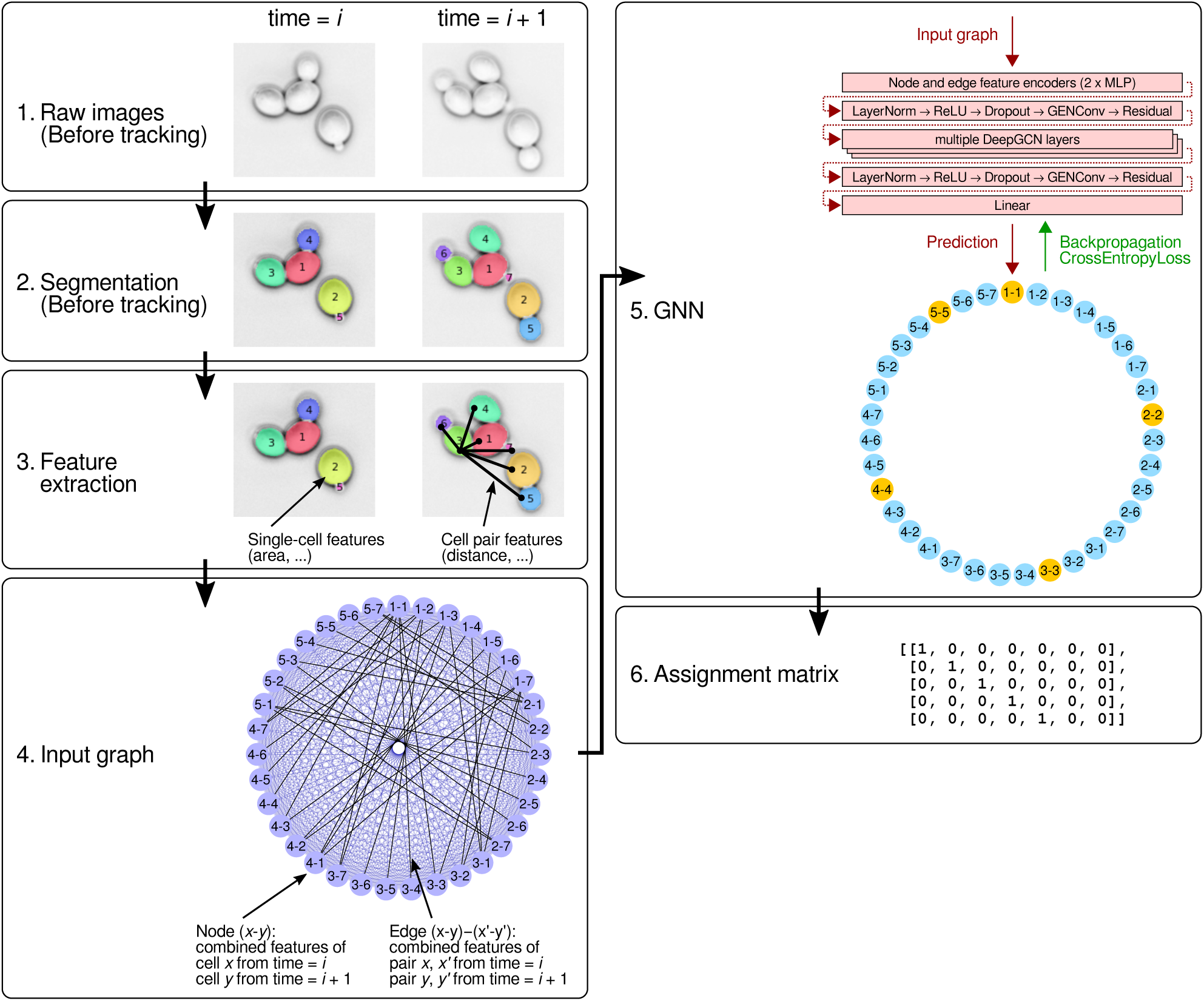
Schematic of the tracking algorithm LYN-track (Steps 3-6). For Steps 1 and 2, we show a field of view with only a few cells for simplicity; however, we tested the tracking algorithm on much larger, densely packed colonies (**Table 1**). In Step 3, single-cell and cell pair features are extracted at both timepoints but are illustrated for one timepoint each for clarity. In Step 4, all possible edges are shown in blue, and edges where both cell pairs are below the distance threshold are in black. In Step 5, the correct assignment nodes for the example shown in Steps 1-3 are highlighted in yellow versus incorrect assignment nodes in blue.

**Table 2:** Single-cell and pair features used by LYN-track. Note that the Fourier descriptors were not found to improve the tracking accuracy and were thus omitted in the final tracking GNN. CoM, center of mass.

| Single-cell features | Pair features |
| --- | --- |
| Area | CoM-to-CoM x-distance |
| Length of major axis (by PCA) | CoM-to-CoM y-distance |
| Length of minor axis (by PCA) | CoM-to-CoM length |
| Geometric mean of principal axes | CoM-to-CoM angle of orientation |
| Angle of orientation | Closest distance |
| Eccentricity |  |
| 10 first Fourier descriptors |  |

To examine information processing through the different message passing layers of the GNN, we visualized node features using t-SNE plots (**Fig. 3**). At the level of the input layer, correct and incorrect assignment nodes were not separated well by t-SNE (**Fig. 3** A). However, at increasing depth in the GNN, specifically, in the first, second, and final GNN message passing layers (**Fig. 3 B-D**), the t-SNE algorithm separated correct and incorrect assignment nodes successively better. These results illustrate how the GNN leverages the graph structure and information propagation through neighboring nodes; the improvements with each layer show the network’s ability to capture and utilize information from cell neighborhoods.

**Figure 3:**
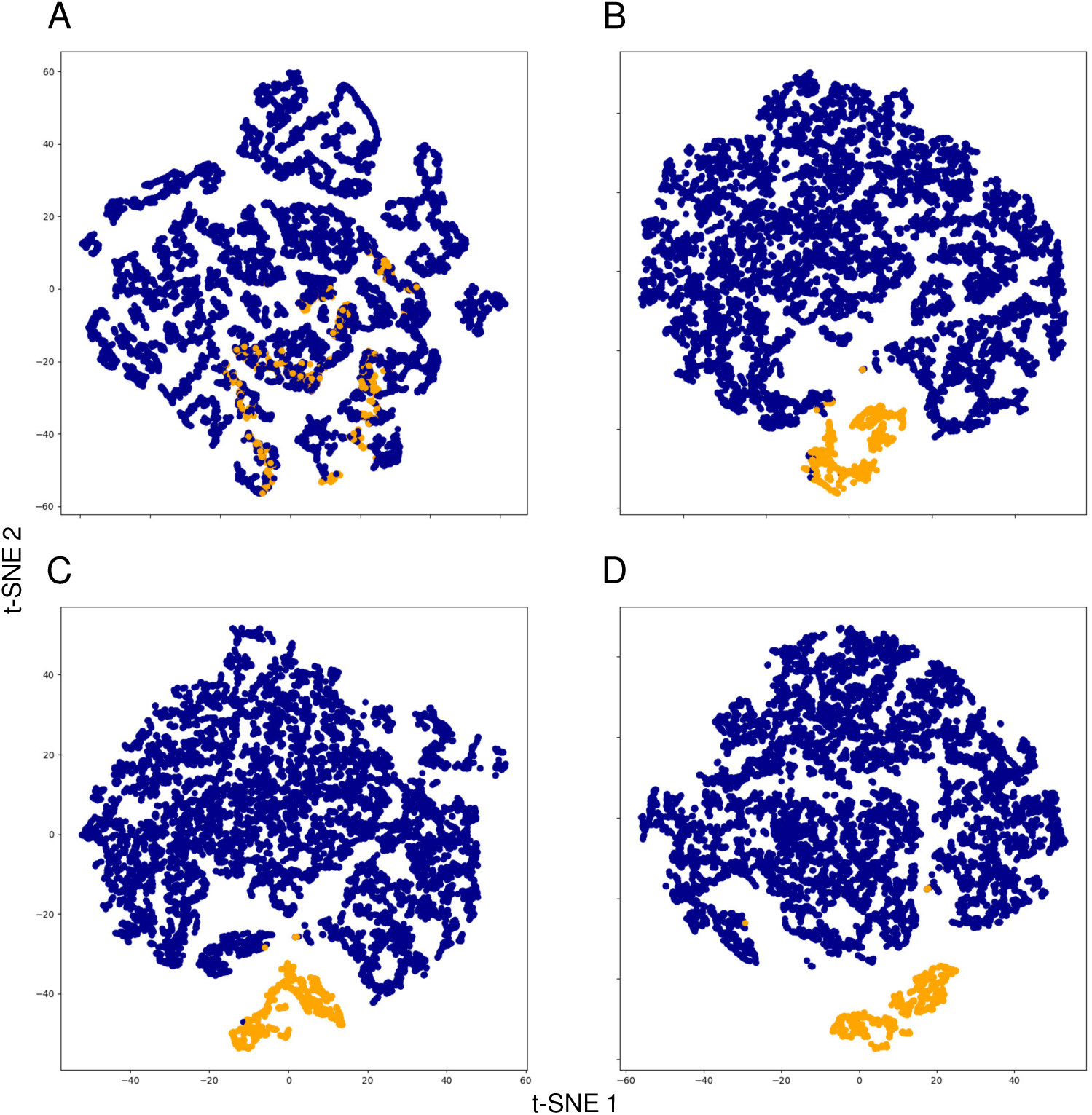
t-SNE plots of node features at different stages of the LYN-track GNN. Yellow and blue nodes represent correct and incorrect assignment nodes, respectively. The GNN was trained on timeframes 2 to 40 from the SJR6-Sc dataset (even frame was the first timepoint and the next (odd) frame the second timepoint). Node representations were obtained for the same pairs of images and visualized all together using t-SNE. Points in the t-SNE plane represent node features in the A) input layer, B) first layer, C) second layer, and D) final layer of the GNN.

### Evaluation of GNN-based tracking algorithm LYN-track

For budding yeast tracking, we used recordings SJR1-Sc to SJR5-Sc for training and recordings SJR6-Sc and SJR7-Sc as validation and test sets, respectively (**Table 1**). To train the algorithm for the more difficult challenge of tracking across larger time intervals, we used the full image sets as well as subsets of images with intervening frames left out to create 10, 15, or 20 min frame intervals. The data from ref.^20^ and from YIT were solely used for testing.

We began by testing an existing generalist tracking method on these budding yeast recordings. Specifically, we evaluated the relatively recent Caliban^36^ algorithm since it bears similarity to our method in so far as it utilizes a graph attention network for encoding neighborhood information but differs in establishing correspondences across time by other (non-GNN) machine learning methods. Caliban was developed for segmenting and tracking mammalian cell nuclei, which divide symmetrically and are well separated. The algorithm requires mother cells to change their cell ID upon budding, which complicates image analysis of budding yeast and necessitated that we adjust the IDs of mother cells in our dataset. We could not run Caliban out of the box (**Methods**). So, we decided to retrain the method using raw images, ground truth segmentation masks, and lineage information from our recordings. After training, Caliban detected only 61% of divisions correctly in the test set (SJR7-Sc), suggesting that more substantial changes are needed to adapt the method to budding yeast.

Considering these results, we benchmarked our method only against algorithms specifically showcased for budding yeast recordings: the state-of-the-art TracX algorithm, which is not based on machine learning and outperforms^20^ previous tracking methods^35,37–40^, as well as the original YeaZ tracking method^18^ (**Table 3**). To evaluate the results, we note that budding yeast recordings are often performed with a 40x or 60x objective and show dozens to hundreds of cells (**Table 1**). Many cell cycle and gene expression events take on the order of minutes to tens of minutes and, thus, timelapse imaging is often performed with frame intervals of ≈3-15 min.^1–7^ Considering these typical recording parameters, accuracies above 99% are desirable so that manual correction followed by computational retracking is only needed infrequently.

**Table 3:** Performance of different tracking methods applied to budding yeast colonies. Values represent F1 scores (mean ± standard error of the mean (SEM)) comparing predicted assignments to correct ground truth assignments. Bold formatting highlights the highest mean score in each row.

| Recording | Frame inter-<br>val [min] | YeaZ | TracX | LYN |
| --- | --- | --- | --- | --- |
| SJR6-Sc (validation set) | 5 | $0.96 \pm 0.01$ | $0.997 \pm 0.002$ | **$0.998 \pm 0.002$** |
| | 10 | $0.93 \pm 0.02$ | $0.98 \pm 0.02$ | **$0.992 \pm 0.003$** |
| | 15 | $0.89 \pm 0.03$ | $0.95 \pm 0.02$ | **$0.982 \pm 0.006$** |
| | 20 | $0.85 \pm 0.04$ | $0.90 \pm 0.04$ | **$0.965 \pm 0.008$** |
| SJR7-Sc (test set) | 5 | $0.992 \pm 0.001$ | $0.998 \pm 0.002$ | **$0.999 \pm 0.001$** |
| | 10 | $0.992 \pm 0.003$ | $0.992 \pm 0.002$ | **$0.999 \pm 0.002$** |
| | 15 | $0.99 \pm 0.02$ | $0.98 \pm 0.02$ | **$0.996 \pm 0.003$** |
| | 20 | $0.96 \pm 0.02$ | $0.96 \pm 0.02$ | **$0.991 \pm 0.004$** |
| TTS-SC7-Sc<sup>20</sup> | 5 | $0.9997 \pm 0.0009$ | $0.998 \pm 0.001$ | **$1.000 \pm 0.001$** |
| | 10 | $0.993 \pm 0.002$ | $0.997 \pm 0.002$ | **$0.998 \pm 0.004$** |
| | 15 | $0.96 \pm 0.03$ | $0.98 \pm 0.02$ | **$0.984 \pm 0.006$** |
| | 20 | $0.90 \pm 0.06$ | **$0.95 \pm 0.04$** | $0.94 \pm 0.01$ |

LYN-track consistently showed high tracking accuracies, especially for 5 min or 10 min imaging intervals. (We gauge the tracking accuracy by the F1 score, in which we score correct or incorrect predicted assignments only with respect to correct ground-truth assignments.) A drop-off in tracking accuracy could be observed beginning with a frame interval of 15 min in recording SJR6-Sc (validation set) and the external TTS-SC7-Sc dataset, both of which have over 300 cells in the last frames. (Note that it is often undesirable or infeasible to decrease the frame interval experimentally (**Discussion**).) TracX showed comparable results on the TTS-SC7-Sc recording, which was supplied with the TracX publication. However, its accuracy was lower beyond the 10 min frame interval for our recordings, highlighting lower generalizability, which cannot straightforwardly be improved by retraining with additional ground-truth annotated images since TracX is not based on machine learning. YeaZ generally performed worse than both other methods. We tested TracX and LYN-track also on the YIT data. Here, both methods gave roughly similar, good results (**Supplementary Table 1**) – but the high frame rate as well as the very small number of frames and cell divisions in these recordings only allowed us to conclude that neither LYN-track nor TracX was extremely sensitive to the underlying dataset.

To evaluate the benefit of our deep learning approach for tracking, we took the cell features that were used as node features for LYN-track (**Table 2**) and computed L2 distances between pairs of cells at consecutive time points. We optimized the weights by minimizing tracking errors, and tested the predictions on SJR6-Sc and SJR7-Sc. The results were worse than LYN-track (**Supplementary Table 2**), underscoring the power of our deep learning approach.

In order to test the generalizability of LYN-track to other cell types, we trained our model on the fission yeast datasets SH1-Sp–SH5-Sp, choosing subsets of images to create three different frame intervals, 5 min, 10 min, and 15 min. To account for the symmetric nature of fission yeast cell divisions, we assigned two new labels to any two new post-division daughters in the datasets and incorporated this constraint into the linear optimization in Step 6 (**Fig. 2**). We then tested the model on the SM1-Sp–SM4-Sp fission yeast colonies that came from another laboratory (**Table 4**). LYN-track performed almost perfectly, and TracX had a slightly higher error rate. These results suggest that tracking fission yeast cells may be overall easier given the distinct length scales associated with rod shapes.

**Table 4:**
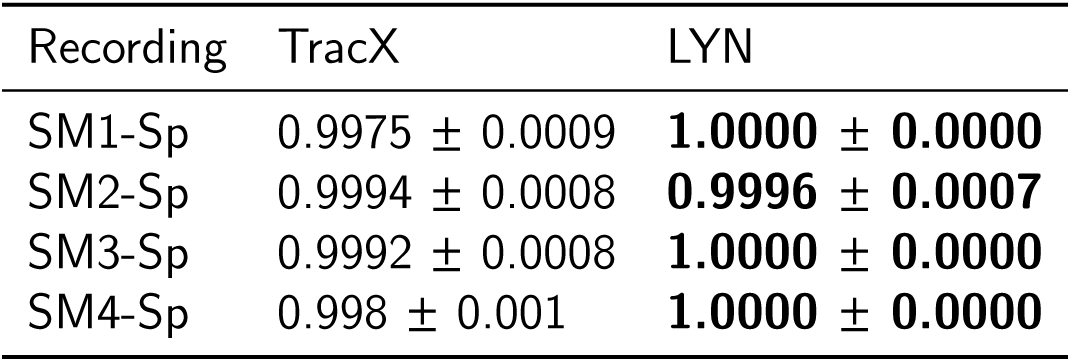
Performance of different tracking methods applied to fission yeast colonies. Values represent F1 scores (mean ± standard error of the mean (SEM)) comparing predicted assignments to correct ground-truth assignments. Bold formatting highlights the best result in each row.

| Recording | TracX | LYN |
| --- | --- | --- |
| SM1-Sp | 0.9975 ± 0.0009 | **1.0000 ± 0.0000** |
| SM2-Sp | 0.9994 ± 0.0008 | **0.9996 ± 0.0007** |
| SM3-Sp | 0.9992 ± 0.0008 | **1.0000 ± 0.0000** |
| SM4-Sp | 0.998 ± 0.001 | **1.0000 ± 0.0000** |

Although designed for tracking in dense colonies, we also tested LYN-track on fluorescently marked mammalian cell nuclei^14^. Nuclei in these datasets divide roughly symmetrically, and are spread unevenly and with large gaps over the field of view due to the comparatively low density of cells and the variable location of each nucleus inside the cell. LYN-track performed well on the higher-magnification dataset (Fluo-N2DH-GOWT) with more uniformly spread nuclei and cells, while it performed poorly on the low-magnification dataset (Fluo-N2DL-HeLa) with unevenly spread nuclei (**Supplementary Table 3**). In the latter case, the small size of the nuclei and the resulting dearth of cell shape features, in addition to the multitude of length scales associated with the uneven spread, likely make such images not suitable for LYN-track. In contrast, when yeast cells divide, the progeny cells touch each other, defining natural neighbors.

### Overview of FC-NN-based lineage tracing algorithm LYN-trace

Lineage tracing is relatively straightforward in budding yeast when bud necks are fluorescently labeled. To automate the assignment of a mother cell to a new bud in this case, we devised the LYN-trace-fluo algorithm as follows: Once a bud appears, for a number of following time points (8 by default), the pipeline blurs the image, retrieves the highest fluorescence peak along the contour of the new bud, and assigns the cell nearest to the fluorescence peak to be a candidate mother cell. The final mother cell prediction is made by majority vote over the potential mother cells identified at each timepoint.

For the more common and challenging case where a bud neck marker is not available, the LYN-trace algorithm (**Fig. 4**) first selects the closest cells within a certain distance of a new small cell as candidate mother cells (default: closest four cells within a 12 pixel contour-to-contour distance). For each pair of bud and candidate mother, features related to their distance, growth of the bud relative to the candidate, relative orientation, and relative position are extracted from a specific number of frames after the appearance of the new bud (default: 8 frames, which corresponds to approximately half of a cell cycle in glucose medium with 5 min frame intervals), providing a rich, dynamic picture of the budding process. We devised the features (**Table 5**) to capture biological preferences for the location, shape, and growth of the bud. We initially tested 21 such features (**Supplementary Table 4**), which were then pared by eliminating the least important feature iteratively (**Fig. 5**, **Methods**). This strategic pruning resulted in an optimal set of 15 highly impactful features (**Table 5**), which maintain accuracy but reduce model complexity. Finally, these features serve as input to a fully connected deep neural network (FC-NN), which predicts which of the candidates is the most likely mother cell.

**Figure 4:**
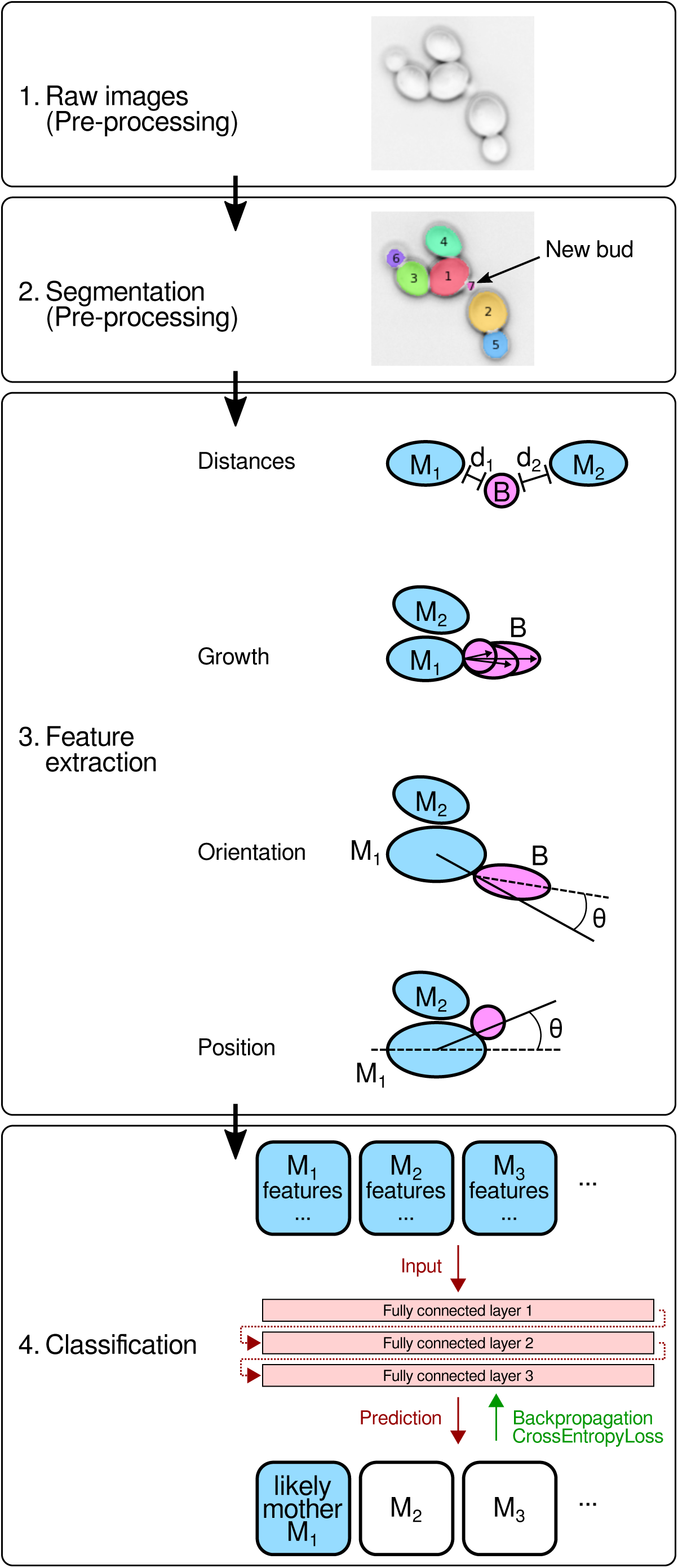
Schematic of LYN-trace (Steps 3-4). Step 3: Black dashed line represents the major axis of a cell. M_i_: i’th candidate mother cell, B: bud.

**Figure 5:**
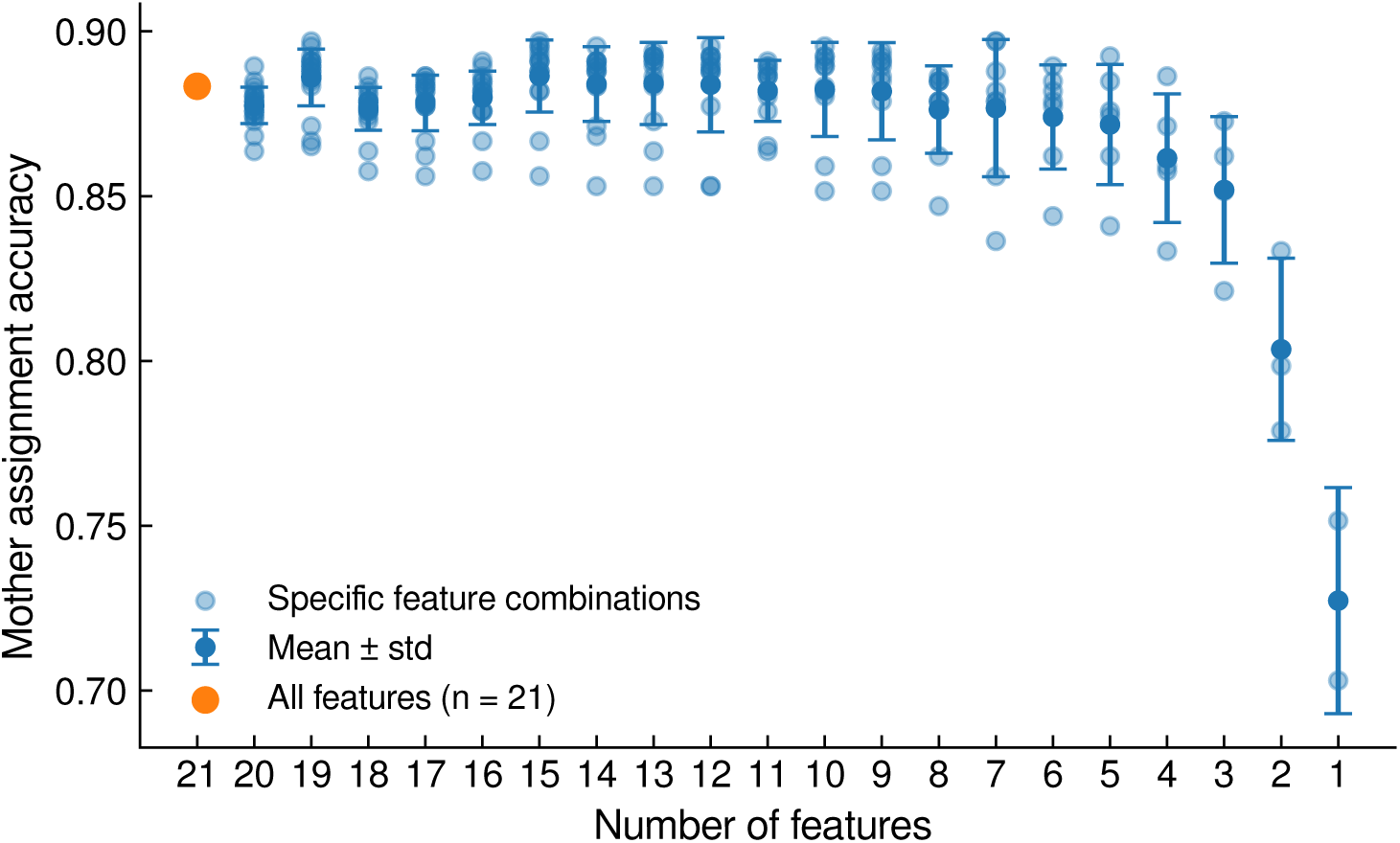
Mother assignment accuracy as a function of the number of features retained after each round of feature elimination. Light blue circles indicate individual candidate feature subsets, dark blue circles indicate the mean, error bars indicate the STD, and the orange marker indicates the baseline model using all 21 features. The highest accuracy is achieved with 15 features. Intermediate subset sizes (≈8-14 features) yielded similar accuracies, whereas smaller subsets (<5 features) showed a marked drop in performance and increased dependence on specific features.

**Table 5:** Features used for lineage tracing. All quantities that require multiple timepoints for calculation, e.g., standard deviation, maximum, minimum, or speed, are computed using *N* frames (default: *N* = 8) following budding. ‘Speed’ is computed by the slope of a linear regression fit. The orientation of the bud is represented by the angle between i) the major axis of the bud and ii) the line from the CoM of the candidate mother cell to the closest point on its contour to the bud. The position angle of the bud along the contour of a candidate mother cell is specified by the angle between i) the major axis of the candidate mother cell and ii) the line between the CoM of the candidate mother cell and the nearest point on the candidate mother contour to the bud.

|  |
| --- |
| Features related to the distance between bud and candidate mother |
| Distance between contours of bud and candidate mother at time of budding |
| Standard deviation of distances between contours |
| Minimum distance between contours of bud and candidate mother |
| Maximum distance between contours of bud and candidate mother |
| Range of distances between contours of bud and candidate mother |
| Features related to the growth of the bud |
| Speed with which bud's CoM moves away from candidate mother contour |
| Speed with which bud's farthest point from candidate mother moves away |
| Features related to the orientation of the bud |
| Bud orientation $N$ frames after budding |
| Change in bud orientation $N$ frames after budding compared to time of budding |
| Minimum of bud orientation |
| Speed of bud orientation change |
| Features related to the position of the bud along the contour of the candidate mother |
| Change in the position angle $N$ frames after budding with respect to at time of budding |
| Standard deviation of position angle |
| Minimum of position angles |
| Maximum of position angles |

### Evaluation of FC-NN-based lineage tracing algorithm LYN-trace

For training, validation, and testing we used a split of 64%, 16%, and 20%, respectively, of all budding events in the SJR recordings (except SJR5-Sc, which lacked bud neck marker images and thus ground truth lineage annotations) (**Table 1**) and 5-fold cross validation. (In comparison, for tracking, the high degree of correlation across time compelled us to separate training, validation, and testing datasets by recording.) We compare LYN-trace to i) the current approach of assigning the nearest cell as the mother cell^21^, ii) LYN-trace-fluo, which can only be used when there is a fluorescent bud neck marker, and LYN-trace with the iii) Random Forest or iv) XGBoost classifier replacing the fully connected neural network (**Table 6**).

**Table 6:** Comparison of the assignment accuracy (correct vs. incorrect) for lineage tracing methods (mean ± standard deviation (STD)). SJR* represents all SJR recordings except SJR5-Sc. Decimals truncated to significant figures. Bold formatting highlights the best result in each row when ignoring LYN-trace-fluo, which has information about the bud neck available.

| Dataset | Nearest cell | LYN-trace -fluo | LYN-trace (Random Forest) | LYN-trace (XGBoost) | LYN-trace |
| --- | --- | --- | --- | --- | --- |
| SJR* (validation set) | | | $0.85 \pm 0.02$ | $0.89 \pm 0.01$ | **$0.90 \pm 0.03$** |
| SJR* (test set) | 0.65 | 1.00 | $0.88 \pm 0.01$ | $0.90 \pm 0.01$ | **$0.910 \pm 0.002$** |
| TTS-SC7-Sc<sup>20</sup> | 0.62 | 0.94 | $0.84 \pm 0.03$ | $0.85 \pm 0.02$ | **$0.864 \pm 0.004$** |

We found LYN-trace to perform substantially better than the nearest-cell assignment, reducing errors by about two thirds. Performance on the external TTS-SC7-Sc recording was slightly lower (although substantially better than nearest-cell assignment), suggesting good generalizability and that training on similar recordings achieves the highest attainable accuracy. Performance with XGBoost was almost as high as with the FC-NN. Our data further show that fluorescent bud neck-based mother assignments can be automated with LYN-trace-fluo to achieve outstanding accuracy.

In order to test whether a multi-layer network was essential for LYN-trace, we retested our method with one of the two hidden layers removed, that is, leaving one hidden layer. The results were substantially worse for the SJR* validation and test sets, highlighting the benefit of the two layers (**Supplementary Table 5**).

## Discussion

Frame-to-frame tracking is highly sensitive to errors because erroneous assignments accumulate and propagate. At the same time, tracking budding yeast cells is highly challenging because cells are densely packed, move due to growth of the colony, change in number, and are poor in distinguishing features. With LYN-track, we leverage geometric features of cells and their neighborhoods to maintain very high tracking accuracies up to 10-15 min intervals between images. Imaging large, rapidly growing colonies in glucose medium with higher frame intervals, that is, 20 min and above, is uncommon^1–7^ and challenging to track by the human eye. (For comparison, under these conditions, a colony doubles in about 90 min.) We do not regard the utility of LYN-track to be in tracking cells in recordings where the user cannot verify the results, e.g., with 20 min intervals and above in glucose medium, but rather in reducing the need for manual correction where the user can still identify the correct tracks.

In light of these results, one may consider simply increasing the frame rate to reduce tracking errors. However, this is often undesirable or infeasible. On the one hand, the size of the raw datasets increases proportionally with the number of images taken, increasing demands on storage and throughput in downstream processing. Furthermore, brightfield imaging (for segmentation and tracking) and fluorescence imaging (for measurements) can often not be uncoupled in microscopy acquisition software, leading to higher phototoxicity merely to achieve higher frame rates for tracking^41–43^.

In this work, we have split up tracking (LYN-track) and lineage tracing (LYN-trace) into two steps. In LYN-track, the same cell is identified in two different images, and new cells are assigned a new ID. In LYN-trace, the mother to a new (daughter) cell is identified. It may be advantageous to merge these algorithms into one, although the ideas presented here do not lend themselves to a combined algorithm easily.

Note that not all tracking tasks share the challenges of tracking budding yeast. Our results indicate that already fission yeast, which is rod-shaped, may provide sufficient shape features to be tracked nearly perfectly (**Table 4**). Blast cells often have highly distinct shapes, and mammalian cell nuclei are generally well separated. Frame-to-frame tracking is not required when tracking body parts of flexible animals^44,45^ or individual animals in a colony^46,47^ since anatomy typically does not change over the course of a recording.

Lineage tracing in timelapse microscopy recordings has received relatively less attention. LYN-trace advances beyond nearest-neighbor mother assignment, which clearly faces limitations inside colonies. We sought to leverage patterns of budding that are inherent to yeast biology to better predict which of multiple candidate mothers gives rise to a new bud.

Finally, all our programs are written in Python, which integrates effectively with other deep learning techniques, such as those used for image segmentation. The overall improvements represented by our methods can be expected to simplify and speed up high-throughput, high-content microscopy.

## Methods

### LYN-track

#### Parameters

Whether an edge is placed in the input graph between two pairs of cells is determined by a parameter which sets the maximum distance between each pair of cells in the same frame (default: 8 pixels).

#### Feature extraction

The features described in **Table 2** are computed using standard numpy and OpenCV (cv2) functions.

#### Cell graph

Each node in the input graph represents the concatenated features of a pair of cells (*x, y*), where *x* is in one time frame and *y* in the next. Edges in the input graph connect two nodes and thus two pairs from the same frame, *x*, *x*^′^ as well as *y*, *y*^′^, and are associated with the concatenated features of the *x*-*x*^′^ and *y*-*y*^′^ pair features. If either of the pairs of cells in the same frame, i.e., *x*-*x*^′^ or *y*-*y*^′^, is further than a Euclidean distance threshold, the edge is not placed. In other words, two nodes (*x*, *y*) and (*x*^′^, *y*^′^) are connected by an edge if max(dist(*x, x*^′^), dist(*y, y*^′^)) is smaller than the distance threshold, where the distance between two cells is measured by the closest points along their contours. We chose 8 pixels as the default distance threshold, which translates to 0.9 *µ*m in our recordings, which is roughly half of the radius of a budding yeast cell.

#### GNN architecture

As illustrated in **Fig. 2**, our model architecture begins with the embedding of the node and edge features of the input graph in two Multi-Layer Perceptrons (MLPs) with ReLU^48^ activation. These layers are followed by a series of convolutions via multiple DeepGCN^49^ layers to perform message passing. Each DeepGCN layer includes a LayerNorm^50^, ReLU^48^ activation, dropout layer^51,52^, GENConv layer^53^, and a residual connection. The GENConv layer processes messages from node and edge features using the following equation:

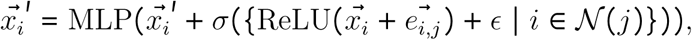

where 𝒩 (*j*) denotes indices of nodes connected to node *j*, *σ* is the softmax function, and *ϵ* is a small learnable constant. The final layer of our model maps the embedded node channels to scores, which are then used in a linear cost minimization algorithm for cell assignment.

#### Data preparation and training

To ensure consistency across varying microscope resolutions and zoom levels, and in cases where metadata is unavailable, all cells in a movie are scaled such that their average area has a fixed value. This standardization is important for model robustness. In addition, we observed that excluding the initial frames, which typically contain fewer cells, from the training set improved model training. Thus, frames with fewer than 20 cells were omitted from the training set although the model remains capable of tracking such scenarios.

#### Assignment matrix

The assignment matrix has *n*_1_ rows and *n*_2_ columns, where *n*_1_ and *n*_2_ are the number of cells in consecutive images. This matrix, illustrated in **Fig. 2**, indicates whether a cell in the first image corresponds to a cell in the second image. The predicted output graph from the GNN contains node weights that after softmax classification can be reshaped into a score matrix, which represents the GNN’s confidence in tracking connections. We employ the linear sum assignment method using the Jonker-Volgenant algorithm in the scipy.optimize package to optimize the total score over all possible matchings resulting in the assignment matrix.

#### Adaptation for fission yeast

The main difference in tracking budding versus fission yeast cells is that the latter divide by growing a septum roughly in the middle of the cell, and splitting the mother cell in two daughter cells. LYN-track is still applicable to fission yeast cells with a minor modification to the linear optimization step to make sure new cells are labeled as such. In the linear optimization step, we ensure that cells that are not tracked to any cell in the previous frame are labeled as new cells by adding the following constraint to the linear optimization problem: If there are *m* cells in the first frame and *n* cells in the second frame and if *m* < *n* holds, we assume that at most *n* − *m* cells have divided, so we have to track at most 2 ∗ *m* − *n* cells to the second frame, and the rest are new cells. The linear optimization problem is solved using the HiGHS solver from the scipy.optimize library.

#### Implementation

LYN-track is implemented in Python and can be used through the YeaZ GUI, which also allows neural network-based segmentation of microscopy images.

#### Caliban

We tried to evaluate Caliban out of the box, i.e., using pretrained network parameters. However, this required the specification of ‘deep cell access tokens’. Unfortunately, despite a number of attempts, we were unable to obtain the tokens, preventing us from pursuing this approach further. Therefore, we turned to retraining Caliban with our data.

### LYN-trace

#### Parameters

LYN-trace-fluo requires a parameter describing the width of the Gaussian kernel to smooth images, which we set by default to 1 pixel. Furthermore, LYN-trace-fluo analyzes by default 8 images after a bud appears to identify the mother cell by majority vote.

LYN-trace analyzes the nearest four cells within a 12 pixel contour-to-contour distance as candidate mothers by default and uses 8 images to compute features.

#### Feature extraction

The features described in **Table 5** are computed using standard numpy and OpenCV (cv2) functions.

#### FC-NN architecture

We define a neural network tailored for the binary classification task of identifying a mother cell from a set of candidate mother cells. The architecture of the neural network is composed of a sequence of fully connected layers with the first layer representing the features of the candidate cells. Three hidden layers apply Exponential Linear Unit (ELU) transformations, chosen for their efficiency in handling gradients and fast training convergence, and include dropout^51,52^ for regularization, reducing overfitting, and batch normalization^50^ to stabilize training and to use nonlinearities effectively. Finally, the output layer is a softmax layer, corresponding to the one-hot encoding of the binary class labels and outputs the confidence of the model.

#### Implementation

LYN-trace is implemented in Python and can be used through a dedicated GUI.

#### Feature elimination

We started with a set of 21 hand-crafted candidate features describing the relationship between a bud and its potential parent cell (**Table 5**, **Supplementary Table 4**). To reduce dimensionality while preserving predictive performance, we applied a greedy backward feature elimination strategy using our neural network classifier as the evaluation model. First, we split the data into a training set (80%) and a held-out test set (20%); the test set was not used to make feature-selection decisions. With the training set, we performed 5-fold cross-validation and, at each elimination step, systematically removed each feature in turn, retrained the network with the remaining features, and computed the mean validation accuracy across the five folds. For each step, we compared these validation accuracies and removed the feature whose exclusion resulted in the highest mean validation accuracy. This procedure was repeated iteratively, reducing the feature set from 21 down to a single feature, while tracking performance at each subset size (**Fig. 5**). We then inspected the cross-validated performance as a function of the number of features. The backward feature elimination analysis showed that the neural network is robust to moderate reductions in input dimensionality but performance degrades once too many features are removed. Across the first several elimination steps, the mean validation accuracy plateaued at high values, in the range of 0.88-0.90, and the highest accuracy was observed for 15 features (0.90 ± 0.03). More aggressive pruning led to a clear drop in validation accuracy, with values falling below 0.88 once only a handful of features were retained, and decreasing sharply when reduced to only two or a single feature. Taken together, these results suggest that a 15-dimensional feature set offers a desirable trade-off between model simplicity and predictive performance, and was thus used as the input for the final model.

### Imaging

The SJR image sets were recorded with a Plan Apo *λ* 60x/1.40 oil objective and a Hamamatsu Orca-Flash4.0 camera. Cells were grown in CellASIC microfluidic chips in standard synthetic complete (SC) medium supplemented with 2% glucose. Images are represented with a 16-bit depth. The diascopic light was generated by Nikon Ti2-E LEDs. The exposure time for the brightfield images was 100 ms. Strain JB207-7C with genotype *MATa ADE2 MYO1-GFP::kanMX HTB2-mCherry::HIS5* was used.

The SH images were recorded at 30^◦^ C on a DeltaVision Elite system (Applied Precision/GE Health-care) equipped with a pco.edge 4.2 sCMOS camera, Olympus 60X/1.42 objective, and an EMBL environmental chamber for temperature control. Cells were grown in Edinburgh minimal medium (EMM, MPBiomedicals, 4110032) and mounted in Y04C microfluidics plates (CellASIC). Cells had either the *cdc2* and *cdc13* genes or the *mad3* gene tagged with GFP and were otherwise wild-type.

The SM images were acquired on a DeltaVision platform (Applied Precision) composed of a customized inverted microscope (IX-71; Olympus), a UPlan Apochromat 100×/1.4 NA oil objective and a color combined unit illuminator (Insight SSI 7; Social Science Insights), using softWoRx v4.1.2 software (Applied Precision). The dataset consisted of mitotically proliferating WT and *pom1*Δ mutant cells imaged with a CoolSNAP HQ2 Photometrics camera and WT mating cells imaged with a 4.2Mpx PrimeBSI sCMOS camera in bin 2×2 mode.

## Declarations

### Ethics approval and consent to participate

Not applicable.

### Consent for publication

Yes.

### Availability of data and material

All images and annotations are available at: https://drive.google.com/file/d/18Lk5eWl6IM_arX7yxIFevVT8jhJaiL8a

[Google drive link to be replaced by Zenodo link upon acceptance of the manuscript.]

All code is available under the MIT license at: https://github.com/rahi-lab/LYN-track-and-trace

### Competing interests

The authors declare no competing interests.

### Funding

Support for FL, VG, ML, RO, GB, FM, MZ, and SJR was provided by SNSF grants CRSK-3 190526, 310030 204938, and CRSK-3 221036 awarded to SJR. SGC and SH were supported by NIH/NIGMS grant R35GM149565 awarded to SH; WL and SGM were supported by ERC Advanced Grant SexYeast awarded to SGM.

### Author contributions

FL, VG, ML, GB, and SJR devised the code. FL, VG, MK, GB, FM, and MZ tested and benchmarked the code. VG, SGC, and WL acquired the data. FL, VG, RO, GB, SGC, FM, and WL labeled data. FL, VG, and SJR wrote the manuscript. SJR, SH, and SGM supervised the work and acquired funding.

## Acknowledgments

We thank Nicole Vadot for early work on the pipeline.

## Supplementary Material

### Supplementary Tables

**Supplementary Table 1:** Performance assessment of TracX and LYN by F1-scores.

| Recording | TracX | LYN-Track |
| --- | --- | --- |
| TTS-YIT-TS1-Sc | 1 | 1 |
| TTS-YIT-TS2-Sc | 1 | 1 |
| TTS-YIT-TS3-Sc | 1 | 0.9972 |
| TTS-YIT-TS4-Sc | 0.9368 | 0.9744 |
| TTS-YIT-TS5-Sc | 0.9891 | 0.9983 |
| TTS-YIT-TS6-Sc | 1 | 1 |
| TTS-YIT-TS7-Sc | 1 | 0.9954 |

**Supplementary Table 2:** Tracking cells by minimizing the L2 norm with the same features used in LYN-Track.

| Dataset name | Subset | F1-score ± STD |
| --- | --- | --- |
| SJR6-Sc (validation set) | 5 min | 0.7 ± 0.2 |
| | 10 min | 0.6 ± 0.3 |
| | 15 min | 0.5 ± 0.3 |
| | 20 min | 0.4 ± 0.3 |
| SJR7-Sc (test set) | 5 min | 0.9 ± 0.1 |
| | 10 min | 0.9 ± 0.2 |
| | 15 min | 0.8 ± 0.2 |
| | 20 min | 0.8 ± 0.2 |

**Supplementary Table 3:** LYN-track applied to mammalian cell nuclei timelapse recordings ^14^.

| Dataset name | Subset | F1-score ± STD |
| --- | --- | --- |
| Fluo-N2DH-GOWT | 01 | 0.99 ± 0.03 |
| Fluo-N2DH-GOWT | 02 | 0.99 ± 0.02 |
| Fluo-N2DL-HeLa | 01 | 0.4 ± 0.3 |
| Fluo-N2DL-HeLa | 02 | 0.3 ± 0.1 |

**Supplementary Table 4:**
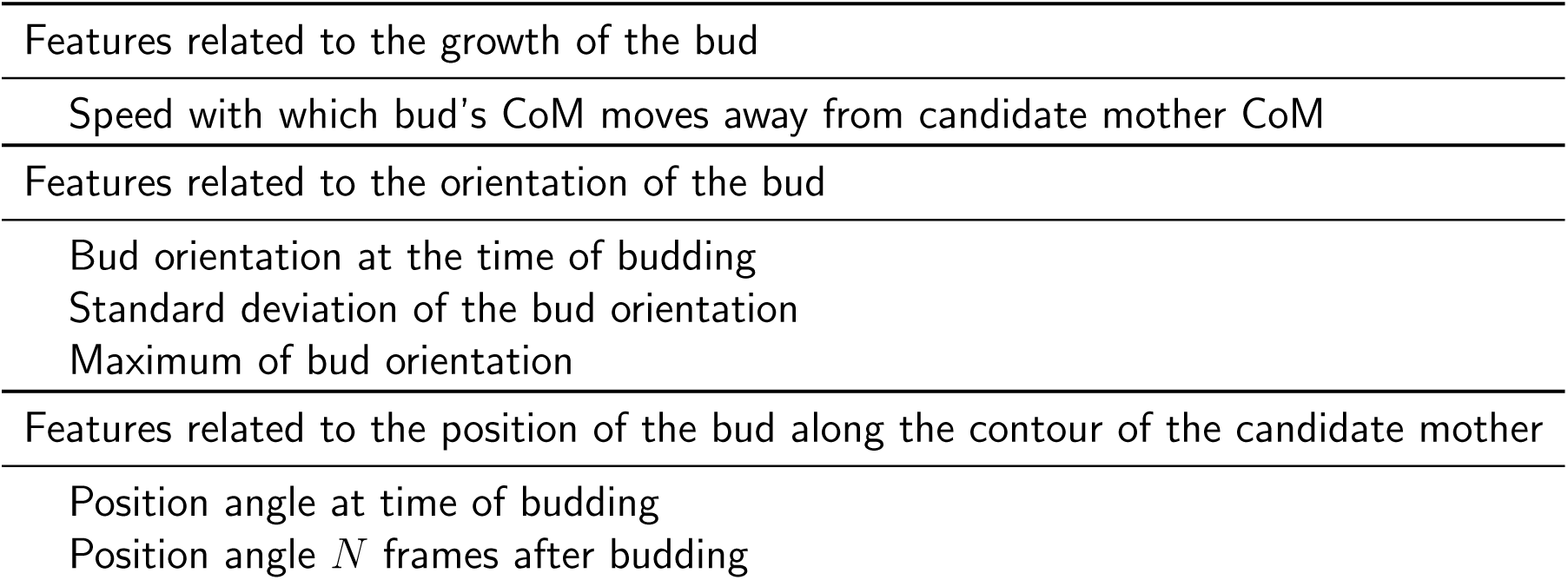
Additional features considered but not used for lineage tracing since they did not increase the performance.

**Supplementary Table 5:** LYN-trace trained and evaluated with only one hidden layer.

| Dataset name | F1-score ± STD |
| --- | --- |
| SJR* (validation set) | 0.73 ± 0.02 |
| SJR* (test set) | 0.70 ± 0.01 |
| TTS-SC7-Sc | 0.837 ± 0.008 |

## Notes

### Competing Interest Statement

The authors have declared no competing interest.

